# Large-scale chemical-genetics of the human gut bacterium *Bacteroides thetaiotaomicron*

**DOI:** 10.1101/573055

**Authors:** Hualan Liu, Morgan N. Price, Hans K. Carlson, Yan Chen, Jayashree Ray, Anthony L. Shiver, Christopher J. Petzold, Kerwyn Casey Huang, Adam P. Arkin, Adam M. Deutschbauer

## Abstract

The genomic catalogue of the human microbiota has expanded dramatically in recent years, and insights derived from human microbiota genomics has vast potential to generate treatments for human diseases. However, predictably harnessing the microbiota for beneficial outcomes is currently limited by our lack of understanding of the physiology of the constituent bacteria. For instance, the functions of most of their genes are not known. Here, we systematically measure mutant phenotypes for genes from the gut commensal *Bacteroides thetaiotaomicron*. Using a barcoded transposon mutant library, we measured the fitness of 4,055 *B. thetaiotaomicron* genes across 492 experiments, including growth on 45 carbon substrates and in the presence of 57 stress-inducing compounds. Our data is in strong agreement with previous studies, and more importantly also uncovers the biological roles of poorly annotated genes. We identified 497 genes with a specific phenotype in only one or a handful of conditions, thus enabling informed predictions of gene function for a subset of these genes. For example, we identified a glycoside hydrolase important for growth on type I rhamnogalacturonan, a DUF4861 protein for glycosaminoglycan utilization, a DUF1080 protein for disaccharide utilization, and a tripartite multidrug resistance system specifically important for bile salt tolerance. Our approach can be applied to other members of the human microbiota to experimentally characterize their genes.

## Introduction

Since the United States National Institutes of Health launched the Human Microbiome Project in 2008 ^1^, our understanding of the human microbiota has advanced significantly. It is now established that the composition and diversity of the gut microbiota play important roles in human health and disease ^2,3^. With these advances, it is now anticipated that manipulation of the human microbiota can be used to treat or manage a number of diseases ^4^. However, these efforts will likely be limited until we have a greater understanding of the activity, physiology, and interactions of the individual bacteria that comprise the human microbiota. One primary bottleneck in our understanding of the human microbiota is that approximately half (40-70%) of the protein-coding genes in the human microbiota do not yet have a predicted molecular function ^5^. Moreover, of the remaining genes, very few have experimentally determined functions; most predictions of gene function are derived by homology to genes characterized in other species from outside the human microbiota, which are often too distant to be accurate ^6,7^.

One of the most well studied constituents of the human microbiota is the Gram-negative gut anaerobe *Bacteroides thetaiotaomicron*, which is prevalent in a substantial fraction of the world’s population ^8^. Pioneering work on the human microbiota revealed the broad range of complex polysaccharides and simple carbohydrates that *B. thetaiotaomicron* can degrade and ferment in the gut ^9-11^. Subsequently, analyses enabled by the full genome sequence of *B. thetaiotaomicron* provided new insights into gene regulation and carbon utilization ^12-14^. It was found that many carbon degradation systems are encoded within Polysaccharide Utilization Loci (PULs), which typically contain a cluster of genes involved in the regulation, transport, and catabolism of various polysaccharides ^15^. The most well studied PUL in *B. thetaiotaomicron* is the starch utilization system (Sus), encoded by *susRABCDEFG* ^16^. SusR is the sensor/regulator, SusCDEF are membrane proteins responsible for starch binding and transport, and SusABG are hydrolytic enzymes. With the minimal requirement of a *susCD* gene pair, genome-wide computational analysis has predicted more than 80 PULs in the *B. thetaiotaomicron* genome ^17^, but the majority of these are experimentally uncharacterized.

Although traditional single-gene genetics and classical enzymology are useful for elucidating protein function, they can be time consuming and labor intensive. As such, it is challenging to apply a “one gene at a time” approach to characterize hundreds to thousands of bacterial genes. Thus, large-scale functional genomic methods are attractive for the systematic characterization of genes without a currently known function ^18^. Methods based on transposon site sequencing (TnSeq) are useful for assaying the importance of thousands of genes in parallel under multiple conditions ^19^, including for human commensals *in vivo*. For example, the transposon insertion sequencing (INSeq) approach was developed in *B. thetaiotaomicron* and revealed genes that were required for colonizing and surviving in mice ^20^. Subsequent application of INSeq to multiple *Bacteroides* species identified species-specific genes important for *in vivo* activity ^21^. The development of large, multi-condition gene phenotype datasets has proven utility for inferring gene function in multiple organisms ^7,22-24^. However, genome-wide fitness assays have yet to be performed systematically across many conditions to identify condition-specific phenotypes in *B. thetaiotaomicron*.

In this work, we describe the use of a barcoded variant of TnSeq (RB-TnSeq) to generate a large genetic dataset for *B. thetaiotaomicron.* In RB-TnSeq, the incorporation of random DNA barcodes simplifies the assessment of strain and gene fitness in a pooled, competitive assay ^25^, and enables the systematic identification of mutant phenotypes across many conditions ^7^, thus massively decreasing the time and effort required for large-scale genetic characterization. Using RB-TnSeq, we performed hundreds of genome-wide fitness assays in *B. thetaiotaomicron,* including in the presence of 113 different compounds such as simple sugars, complex polysaccharides, and antibiotics. This rich dataset provides experimental evidence for many computationally predicted gene functions in *B. thetaiotaomicron*, and also links genes that previously lacked informative annotations to specific processes including polysaccharide degradation, disaccharide catabolism, and bile salt tolerance.

## Results

### A gene-phenotype map of *B. thetaiotaomicron*

To identify an effective transposon delivery vector for *B. thetaiotaomicron* VPI-5482, we first tested the mutagenesis efficiency of hundreds of different vectors in parallel. This approach, which we term “magic pools” ^26^, enabled us to design a *mariner* transposon vector that was suitable for constructing a genome-wide library with minimal insertion biases (Methods). The final mutant library contains 316,025 uniquely barcoded strains with insertions that we could confidently map to a single location in the *B. thetaiotaomicron* genome. These transposon insertions are evenly distributed across the chromosome (Figure 1A), and are equally likely to be on the coding and non-coding strands of the mutated genes (of insertions within protein-coding genes, 49.4% had the drug marker on the coding strand). To accelerate follow-up studies, we also generated an archived collection of individual transposon insertion mutants in 96-well microplate format and mapped the identity of the majority of these strains (A.L.S, A.M.D, K.C.H; in review).

**Figure 1.**
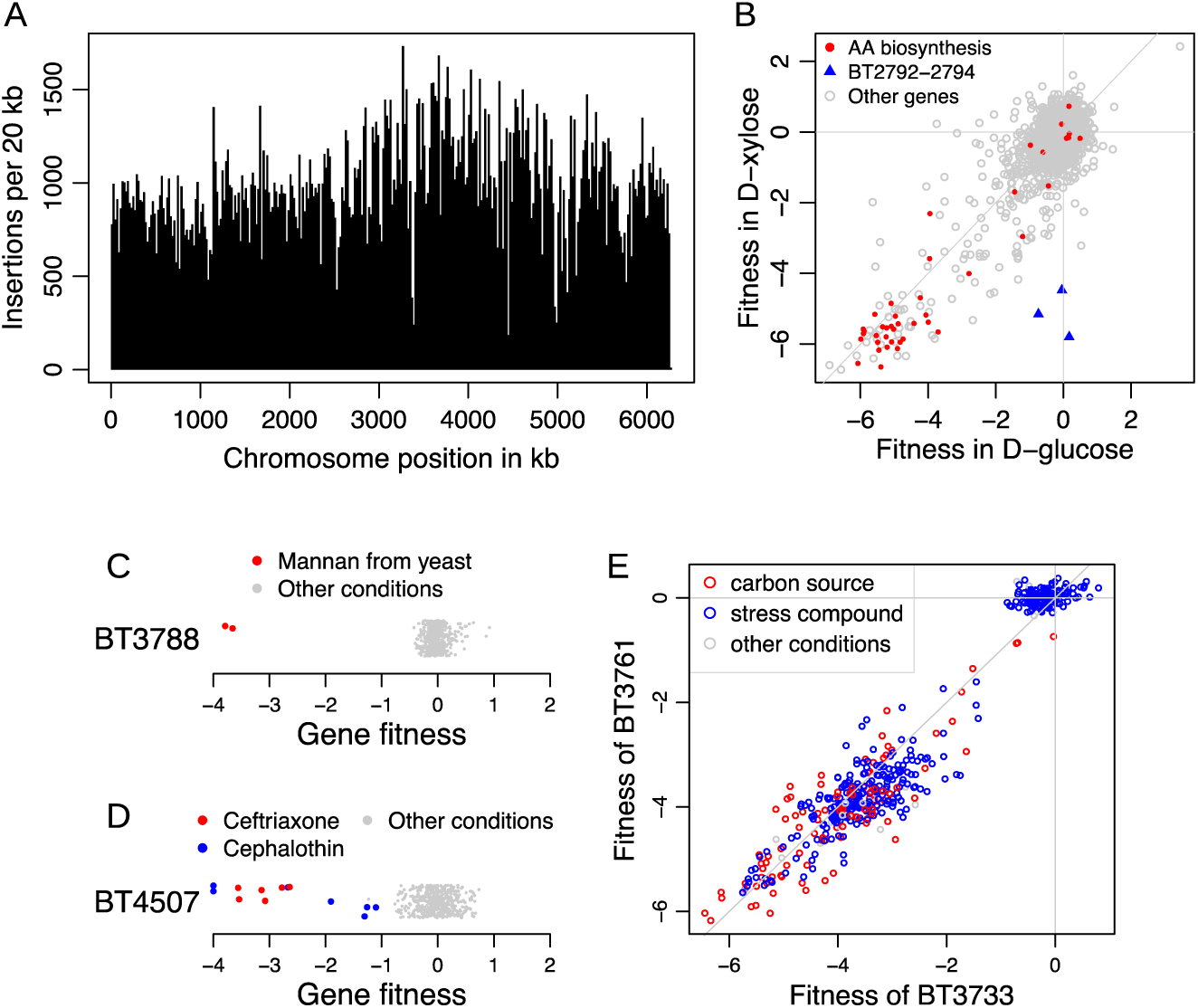
High-throughput genetics in *B. thetaiotaomicron*. (A) Transposon insertions were approximately evenly distributed across the *B. thetaiotaomicron* chromosome. (B) Gene fitness values during growth with glucose or xylose as the sole carbon source were consistent with previous knowledge. The amino acid biosynthesis genes are predicted by TIGR roles ^81^. (C,D) Gene fitness values across conditions revealed that BT3788 is important for fitness on mannan, and mutants in BT4507 were impaired only under treatment with ceftriaxone and cephalothin. The *y*-axis is random and fitness values less than −4 are shown at −4. Negative values mean that the mutant is less fit than the typical mutant. The conditions in which these genes have a specific phenotype are marked. (E) Comparison of gene fitness values for BT3733 and BT3761 across all experiments. Because BT3733 encodes an enzyme in arginine biosynthesis, the high cofitness suggests that BT3761 is also involved in arginine biosynthesis.

To validate the utility of the *B. thetaiotaomicron* mutant library for genome-wide fitness assays, we grew the pooled library in a defined growth medium with either glucose or xylose as the sole carbon source. For each experiment, we measured the abundance of the DNA barcodes by sequencing (BarSeq) before and after growth selection, and we defined the fitness for each gene as the log_2_ change in abundance of mutants of the gene (Figure 1B). Negative and positive values mean that the mutants in that gene were less and more fit, respectively compared to the average strain. Many genes that are predicted to be involved in amino acid biosynthesis were found to be important for fitness with either carbon source, because the defined media lacked most of the amino acids. Genes from the predicted xylose utilization pathway, BT0792 (xylulose kinase), BT0793 (xylose isomerase) and BT0794 (xylose transporter), were important for growth on xylose only (Figure 1B). This simple example illustrates the biological consistency of fitness assays using our mutant library.

The primary advantage of a barcoded transposon mutant library is that each genome-wide fitness assay only requires quantification of the DNA barcodes with BarSeq, which is extremely scalable ^25^. This feature of our method allowed us to perform hundreds of genome-wide fitness assays to create a large gene-phenotype compendium for *B. thetaiotaomicron*. Many of these experiments were assays in defined media (Varel-Bryant (VB)) with a single carbon source for the bacteria to consume. In addition, we performed hundreds of competitive growth experiments with an inhibitory but sublethal concentration of a stressor such as an antibiotic, biocide, or bile salt. Overall, we performed 492 genome-wide fitness assays that passed our metrics for internal consistency ^25^ (Methods). All analyses in this manuscript are derived from these 492 experiments, which represent fitness data in the presence of 113 unique compounds. All data is available online for interactive analyses at the Fitness Browser (http://fit.genomics.lbl.gov).

We successfully measured fitness for 4,055 protein-coding genes in each of these experiments, with the fitness for the median gene calculated from 24 independent transposon insertions. Among the genes with no fitness data, we estimate that 378 of them are likely essential for viability in the Brain Heart Infusion Supplemented (BHIS) media we used to select the mutants (Supplementary Table 1). Some non-essential genes may not have fitness data because they are too short, repetitive, or their mutants are at a low abundance because they are nearly essential for viability.

Across the entire dataset, we identified 497 different genes with a specific phenotype (Supplementary Table 2), defined as |fitness| > 1 and statistically significant in an experiment but lacking a phenotype in most other experiments (Methods). A specific phenotype often links a gene to a condition that is directly related to its function ^7^. To illustrate this principle, we present examples of specific phenotypes for BT3788 and BT4507. BT3788 is a SusC homolog located in the PUL for mannan utilization ^27^. Across all 45 different carbon sources we profiled, BT3788 was only important for fitness on mannan (Figure 1C). BT4507 encodes a class A serine β-lactamase ^28^, and we found that mutants in this gene were only impaired in the presence of two β-lactam antibiotics, ceftriaxone and cephalothin, but not in any of the other conditions that we profiled, including dozens of other antibiotics (Figure 1D).

A second approach for inferring gene function from large-scale genetics data is cofitness, where a pair of genes from the same organism have very similar fitness profiles across all conditions ^7^. To illustrate the utility of cofitness to infer gene function in *B. thetaiotaomicron*, we investigated genes important for arginine biosynthesis. The predicted operon BT3758-3762 encodes several homologs to genes in the *Escherichia coli* arginine biosynthesis pathway ^29^, except that BT3761 is not closely related to an *E. coli* enzyme (by BLAST), but rather shares homology to Cabys_1732 from *Caldithrix abyssi*. These two proteins share 58% identity with 88% sequence coverage. Cabys_1732 is annotated as a newly characterized isoform of N-acetylglutamate synthase ^30^. From our data, we found that BT3761 has very high cofitness (>0.92) with the other four genes in the same putative operon, and it has the highest cofitness (0.97) with BT3733 (Figure 1E), which is a homolog of *E. coli* argininosuccinate lyase. Based on these data, BT3761 likely encodes an N-acetylglutamate synthase required for arginine biosynthesis. A full list of genes with high cofitness is available in Supplementary Table 3.

Overall, we identified a functional association (either a specific phenotype or cofitness) for 612 of the 4,055 protein coding genes for which we collected data. Genes with a functional association were more likely to have a specific COG functional category assigned compared to genes with no functional association (58% vs. 38%; *P* < 10^-15^, Fisher’s exact test) ^31^. Nonetheless, we were able to identify a functional association for 256 genes without a specific COG function category. According to the PaperBLAST database ^32^, 193 of these 256 genes have not been previously discussed in the literature, nor have close homologs (at >70% amino acid identity) from other *Bacteroides* species.

### Linking PULs to specific substrates with genetics

To associate the PULs of *B. thetaiotaomicron* to their substrates, we analyzed the fitness data collected by growing the mutant library in defined media with various compounds added as the sole source of carbon. The 45 carbon sources we successfully assayed represent a range of substrate complexity and include 15 monosaccharides and their derivatives, 10 disaccharides, 1 trisaccharide, 2 tetrasaccharides, 3 oligosaccharides (>4 monomers), and 14 complex polysaccharides. The complex polysaccharides we assayed are typically derived from natural sources (for example, starch from potato) and are likely more heterogeneous than the simpler sugars; we note the source of each compound in Supplementary Table 4. Lastly, because we performed all of our genome-wide fitness assays in pools, we cannot rule out the possibility of cross-feeding between strains, with the potential implication that some genes will have no phenotype in our data despite their importance for growth on a particular substrate.

Fitness profiling across a range of substrate complexity provides a broad view of the importance of each PUL component to the catabolism of a complex polysaccharide. As an illustration, we re-examined the canonical Sus system for starch degradation by comparing the phenotypes of these genes during growth on starch, the component polysaccharides of starch (amylopectin and amylose), the oligosaccharide α-cyclodextrin, and the simpler starch break-down products glucose, maltose, maltotetraose, and maltohexaose (Figure 2). The SusCD transporter proteins were important for fitness on all saccharides of length 4 or more, but largely dispensable for growth on maltose and glucose, which agrees with previous findings that the importance of SusD varies depending on the size of the glucose polymers ^33^.

**Figure 2.**
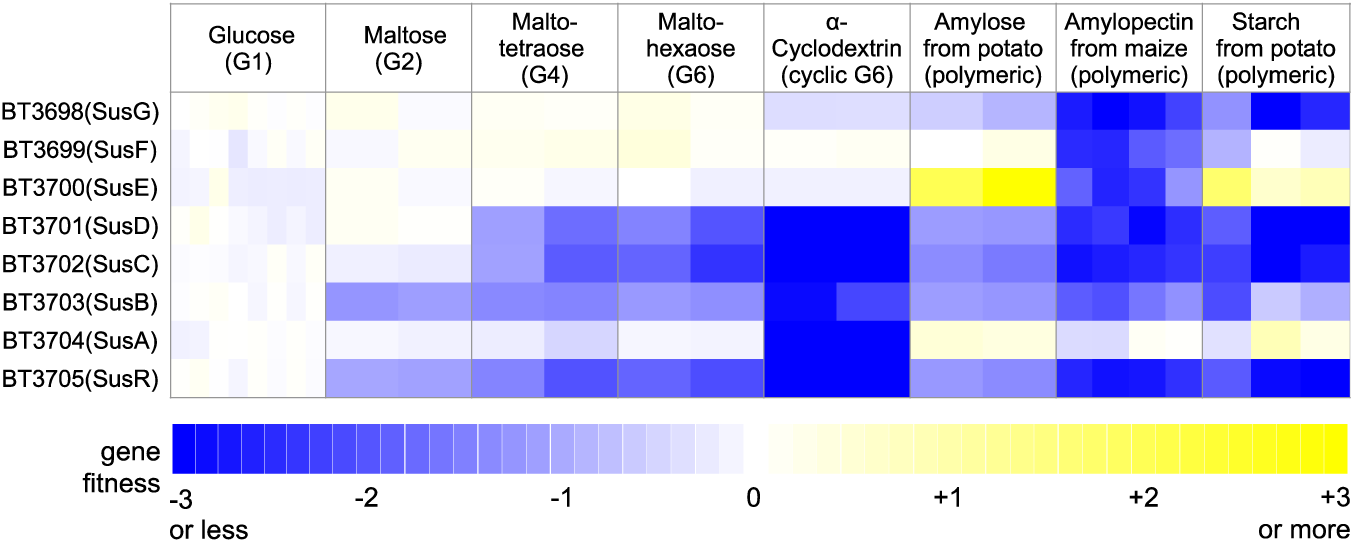
Fitness data for the Sus PUL shows a broad range of importance for substrate catabolism. A heatmap of gene fitness values for the starch utilization system genes during growth on different glucose-containing carbon sources is shown. The data from each replicate experiment is shown separately, for example 6 glucose experiments and 4 amylopectin experiments. The compounds are ordered from left to right by increasing complexity.

The three glycoside hydrolases (GHs), SusA, SusB, and SusG, were variably important across these substrates. SusA was previously characterized biochemically as a neopullulanase that acts to cleave alpha-1,4 linkages in pullulan, a polymer of maltotriose subunits, and was shown to be mildly important for growth on starch ^34^. In our data, SusA is only important for growth on α-cyclodextrin, a cyclic oligosaccharide of 6 glucose monomers linked by α(1-4) glycosidic bonds (Figure 2). SusA homologs in other species are often annotated as cyclomaltodextrinase (an enzyme that linearizes circular oligosaccharides), and homologs of this protein are required for growth on α-cyclodextrin in diverse bacteria including *Caulobacter crescentus* and *Echinocola vietnamensis* KMM 6221 ^7^. These data suggest that SusA also has cyclomaltodextrinase activity, as has previously been proposed ^35^. SusB is a GH97 family member that hydrolyzes multiple linkages, including α(1-4) bonds ^36^. In contrast to a previous study that found only a minimal growth defect of a *susB* mutant on starch ^34^, our results showed that SusB was required for optimal growth on all of these saccharides except glucose, suggesting a key role in hydrolyzing diverse starch derivatives, including the simple maltose disaccharide. SusG is an alpha-amylase localized to the cell surface that breaks down starch into simpler oligosaccharides ^37^, which is supported by our fitness data: SusG was very important for growth on starch and amylopectin, and to a lesser extent on amylose and α-cyclodextrin (Figure 2). Lastly, while the outer membrane proteins SusE and SusF have been demonstrated to bind starch ^38^, their precise function remains unclear. We found that these proteins were only important for growth on amylopectin, suggesting that SusEF preferentially bind this component of starch or are important for stabilization of the SusG-starch complex. Taken together, our re-evaluation of the starch utilization system based on genetics data across multiple substrates provides new insight into this well-studied system.

To globally associate PULs to different carbon substrates, we identified instances where at least one gene in a given PUL had a specific and important phenotype (gene met criteria for specific phenotype and fitness < −1) on a small number of the 45 carbon sources. Based on these criteria, we were able to associate 20 of the 93 PULs (from ^39^) to 32 substrates (Table 1, Figure 3). Some PULs were only important for growth on a single compound, for example PUL85 and heparin, while others were important on multiple substrates (such as the aforementioned Sus locus). Conversely, some compounds could be linked to a single PUL, for example dextran with PUL48, while others like arabinogalactan appear to be catabolized by multiple PULs (Figure 3). Overall, our experimental data is in strong agreement with previous findings for several PULs. We identified specific and important phenotypes for at least 3 genes from multiple PULs on their predicted or known polysaccharides (Table 1, Figure 3): PUL5 on arabinogalactan ^40^; PUL22 on fructans ^41^; PUL48 on dextran ^42^; PUL57 on chondroitin sulfate and hyaluronic acid ^43^; PUL68 on mannan ^27^; PUL75 on pectic glycans ^44^; PUL85 on heparin ^45^; and PUL86 on type I rhamnogalacturonan^44^.

**Figure 3.**
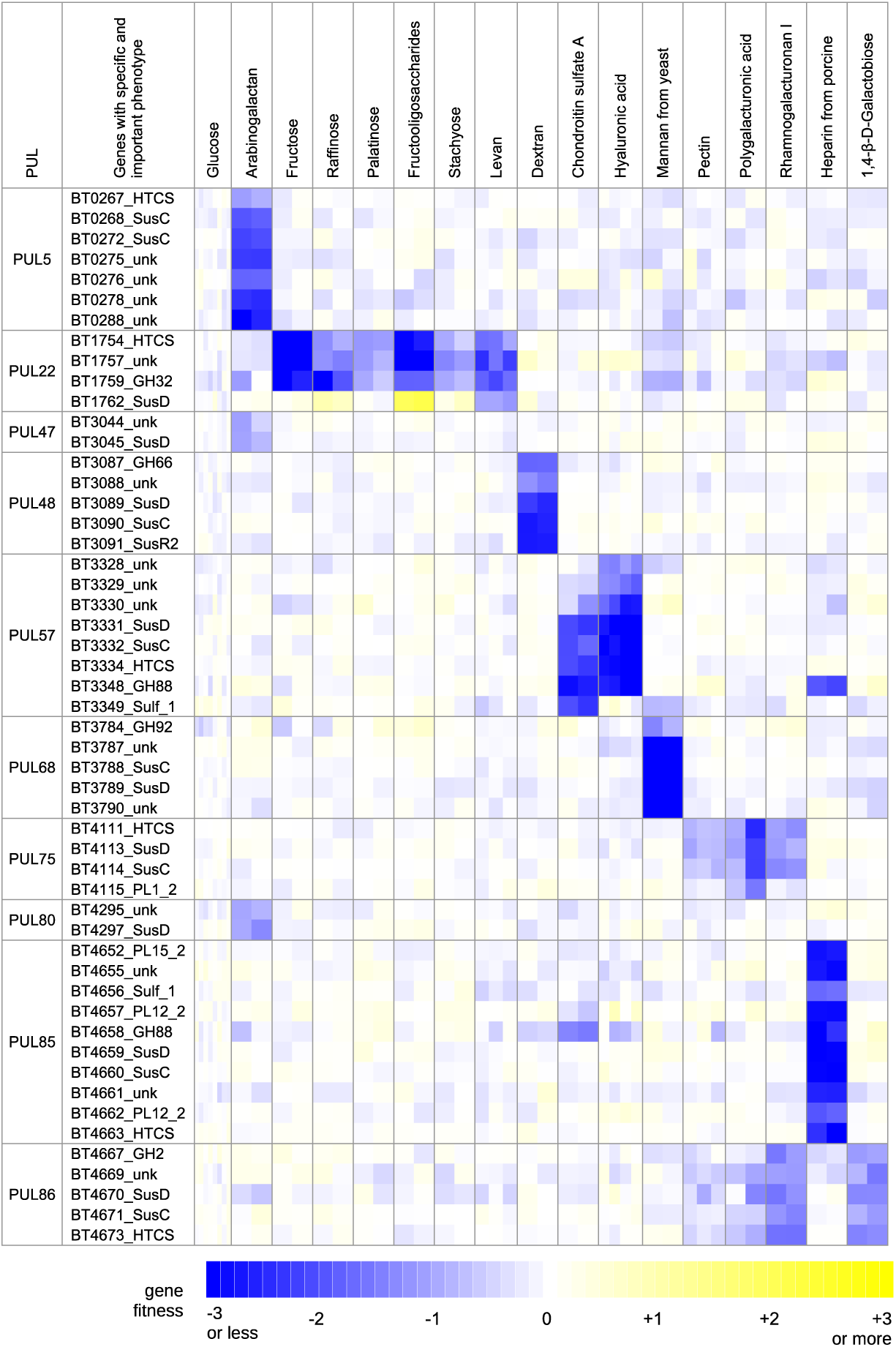
Fitness data associates PULs with substrates. A heatmap of fitness data for select genes with important (fitness < −1) and specific phenotypes during growth on various carbon sources is shown. We present the fitness data for each replicate experiment separately. HTCS: AraC-like hybrid two-component system; unk: unknown; GH: glycoside hydrolase; PL: polysaccharide lyase. The labels indicate gene families and annotations are derived from PULDB ^39^.

**Table 1.** Genes with specific and important phenotypes for 20 PULs on various carbon sources. A PUL was included in the table if at least one of its component genes was specifically important for fitness (fitness < −1) on one or a few of the 45 carbon sources we profiled. ^a^Literature-derived PULs from CAZy database (http://www.cazy.org/PULDB/) ^39^;^b^Proteins in the PUL; ^c^Carbohydrate conditions in which gene(s) from the PUL have a specific and important phenotype.

| PUL<sup>a</sup> | Genes<sup>b</sup> | Carbohydrates<sup>c</sup> |
| --- | --- | --- |
| 5 | BT0262-0291 | Arabinogalactan (BT0267, 0268, 0272, 0275, 0276, 0278, 0288) |
| 7 | BT0348-0356,<br>0358-0369 | D-Arabinose (BT0351, 0355, 0356);<br>L-Arabinose (BT0350, 0351, 0355, 0356) |
| 15 | BT1117-1119 | 1,4-β-D-Galactobiose (BT1118); Galacturonic acid (BT1118, 1119) |
| 22 | BT1754,<br>1757-1763,<br>1765 | Fructose (BT1754, 1757, 1759); Raffinose (BT1754, 1757, 1759);<br>Palatinose (BT1754); Fructooligosaccharides (BT1754, 1757, 1762);<br>Stachyose (BT1754); Levan (BT1754, 1757); |
| 24 | BT1871-1878 | Melibiose (BT1871) |
| 36 | BT2615-2633 | Mannan from yeast (BT2629) |
| 37 | BT2802-2809 | Ribose (BT2802, 2803, 2804, 2809) |
| 47 | BT3043-3049 | Arabinogalactan (BT3044, 3045) |
| 48 | BT3086-3091 | Dextran (BT3087, 3088, 3089, 3090, 3091); Red arabinan (BT3089) |
| 57 | BT3324,<br>3328-3334,<br>3348-335 | Chondroitin sulfate A (BT3330, 3331, 3332, 3334, 3348, 3349);<br>Hyaluronic acid (BT3328, 3329, 3330, 3331, 3332, 3334, 3348);<br>Heparin (BT3348) |
| 59 | BT3461-3507 | Fructooligosaccharides (BT3485); Hyaluronic acid (BT3487) |
| 61 | BT3566-3570 | Laminaribiose (BT3567, 3568, 3569) |
| 66 | BT3698-3705 | Maltose (BT3703, 3705);<br>Maltotetraose (BT3701, 3702, 3703, 3705);<br>Maltohexaose (BT3701, 3702, 3703, 3705);<br>α-Cyclodextrin (BT3701, 3702, 3703, 3704, 3705);<br>Amylose (BT3701, 3702, 3703, 3705);<br>Amylopectin (BT3698, 3699, 3700, 3701, 3702, 3703, 3705);<br>Starch (BT3698, 3701, 3702, 3703, 3705)<br>Polygalacturonic acid (BT3698, 3701, 3702, 3705); |
| 68 | BT3773-3782,<br>3784,<br>3786-3792 | Mannan from yeast (BT3784, 3787, 3788, 3789, 3790) |
| 75 | BT4108-4124 | Pectin (BT4113);<br>Polygalacturonic acid (BT4111, 4113, 4114, 4115);<br>Rhamnogalacturonan I (BT4111, 4113, 4114) |
| 77 | BT4145-4183 | Rhamnogalacturonan I (BT4169) |
| 80 | BT4294-4300 | Galacturonic acid (BT4300); Arabinogalactan (BT4295, 4297) |
| 84 | BT4631-4636 | Arabinogalactan (BT4636) |
| 85 | BT4652-4663,<br>4675 | Heparin (BT4652, 4655, 4656, 4657, 4658, 4659, 4660, 4661, 4662, 4663) |
| 86 | BT4667-4673 | 1,4-β-D-Galactobiose (BT4667, 4669, 4670, 4671, 4673);<br>Rhamnogalacturonan I (BT4667, 4670, 4671, 4673);<br>Polygalacturonic acid (BT4670) |

In addition to confirming the results of these previous studies, we provide the first genetic evidence for the prediction that, besides PUL5, PUL47 and PUL80 are also important for arabinogalactan utilization ^46^. The arabinogalactan phenotypes for genes in PUL47 and PUL80 were mild (fitness values ranged from −0.7 to −1.4 for statistically significant phenotypes), but these mild phenotypes were observed across at least 3 genes in each PUL, and the genes in these clusters do not have clear phenotypes in any of the other conditions that we profiled (Figure 4A,B). These data demonstrate that the efficient breakdown of certain complex polysaccharides like arabinogalactan requires multiple PULs. It was also proposed that PUL65 (BT3674-3687) is involved in arabinogalactan breakdown ^40^, but no genes in this cluster had a strong phenotype in our dataset, including on arabinogalactan (all genes fitness > −0.5).

**Figure 4.**
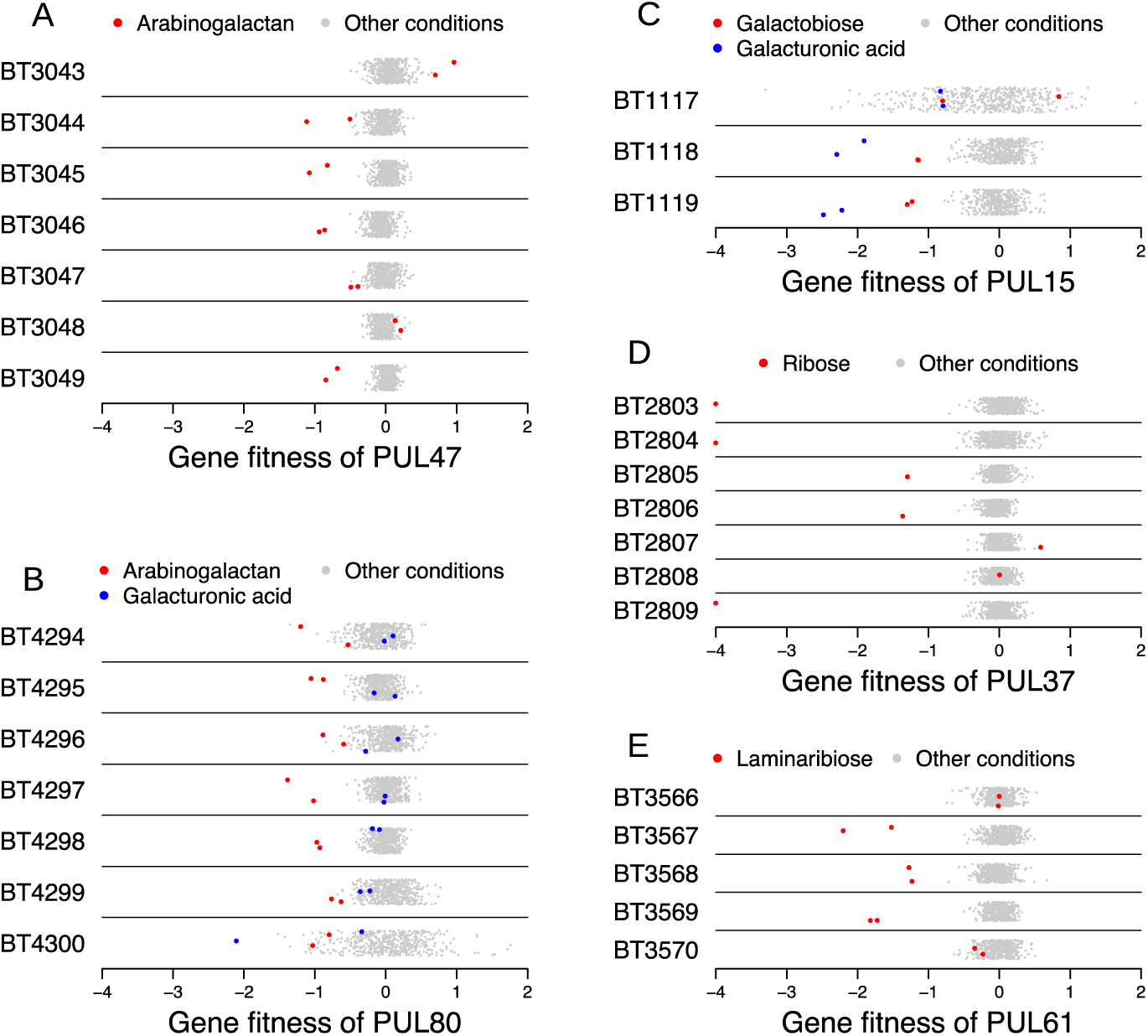
Fitness data for select PULs provides genetic evidence for novel substrates. For the indicated PUL, we show the fitness data for all component genes with certain carbon source conditions highlighted. The y-axis is random, and values less than −4 are shown at −4.

Finally, our analysis was able to link some previously uncharacterized PULs to specific monosaccharides and disaccharides (Figure 4C-E). For example, the SusCD homologs of PUL15 (BT1118-1119) were important for growth on galacturonic acid and 1,4-β-D-galactobiose (Figure 4C). In addition, we found that components of PUL37 were only important for growth on D-ribose (Figure 4D), in particular a hypothetical protein (BT2802), two predicted ribokinases (BT2803-2804), and a sugar transporter (BT2809). The SusCD homologs in this cluster were also mildly important for growth on D-ribose (fitness of BT2805 = −1.3 and fitness of BT2806 = −1.4). Lastly, we found that a putative glucosidase (BT3567), SusD-like protein (BT3568), and SusC-like protein (BT3569) from PUL61 were required for optimal growth on the disaccharide laminaribiose (Figure 4E). While the PULs of *B. thetaiotaomicron* have typically been studied in the context of complex polysaccharides, our data suggests that some of these systems may have evolved to specialize in the catabolism of simpler substrates. In support of this view, PUL37 and PUL61 contain the core SusCD proteins, but do not have homologs of SusEF or hydrolases expected to act on complex substrates. Also, in each instance, the SusCD proteins are important for using simple sugars, suggesting that they bind these substrates on the outer surface of the cell.

### Identification of new polysaccharide degradation genes

Although the degradation pathways for a number of polysaccharides are well described in *B. thetaiotaomicron*, it is likely that additional genes outside of the predicted PULs are also involved in these processes. Indeed, prior work demonstrated that genes important in glycan utilization can be located outside the corresponding PUL(s) ^43,47^. By examining genes with specific and important (fitness < −1) phenotypes, we identified three new genes important for the utilization of these polysaccharides.

The first example is in the metabolism of type I rhamnogalacturonan (RG-I), a polysaccharide containing repeats of a disaccharide of galacturonic acid and rhamnose. A recent study proposed the involvement of five GH28-family glycosyl hydrolases (BT4123, BT4146, BT4149, BT4153, and BT4155) in breaking down RG-I ^44^. In our fitness data, only BT4149 was important for utilizing RG-I, although this phenotype was mild (fitness = −1.3). Rather, we found that another GH28-family member, BT4187, was more important for growth on RG-I (fitness = −2.2) (Figure 5). In further support of the importance of BT4187 in RG-I degradation, the operon containing BT4187 is predicted to be in the same regulon as PUL75 ^13^, which is both important for fitness (Figure 3) and induced during growth on RG-I ^46^. While it is possible that the other hydrolases are functionally redundant, our results demonstrate that the loss of BT4187 alone results in a strong fitness defect during growth on RG-I, suggesting that BT4187 has a unique hydrolase activity.

**Figure 5.**
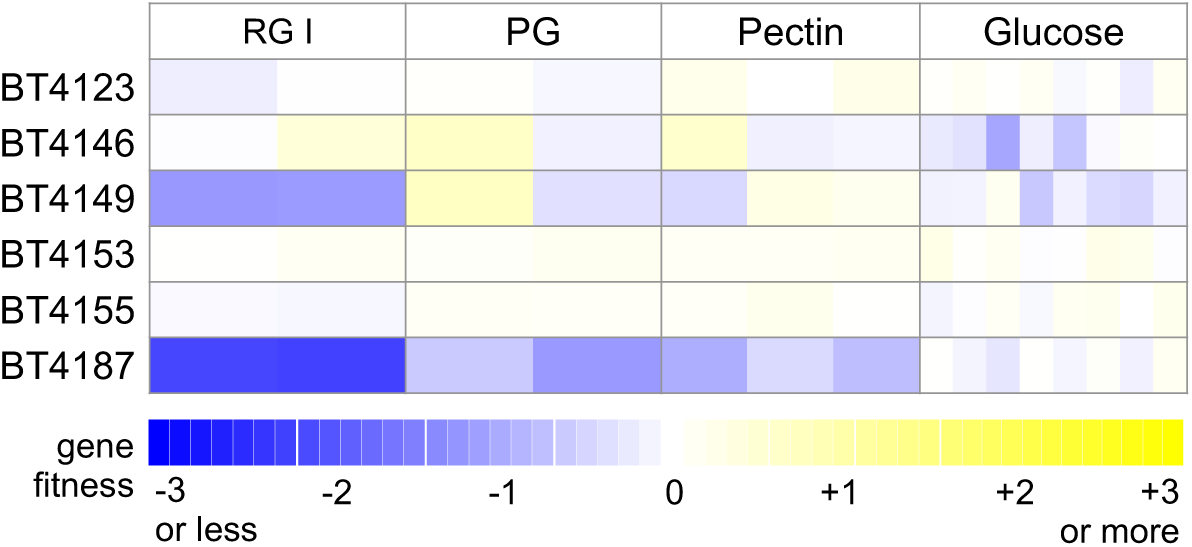
BT4187 is important for type I rhamnogalacturonan utilization. Mutant fitness data for six predicted GH28-family glycosyl hydrolases on type I rhamnogalacturonan (RG-I), polygalacturonic acid (PG), pectin, and glucose. Each replicate experiment is shown separately.

The next two examples are previously unknown genes for utilizing host-derived glycosaminoglycans (GAGs), which are structural components of animal tissues and are composed of repeating disaccharide units of a hexuronic acid linked to an amino sugar. *B. thetaiotaomicron* is capable of using multiple GAGs as nutrients, including chondroitin sulfate (CS), hyaluronic acid (HA), and heparin ^45,48,49^. CS and HA differ by their repeating disaccharide: CS contains sulfated N-acetylgalactosamine and glucuronic acid, and HA contains N-acetylglucosamine and glucuronic acid. Previous studies have proposed a model for CS and HA catabolism by *B. thetaiotaomicron* involving PUL57 and BT4410, a CS lyase ^17,43^. In our experiments, many components of PUL57 were indeed required for optimal growth on CS and HA, with the SusCD homologs (BT3331-3332), glucuronyl hydrolase (BT3348), and hybrid two-component regulator (BT3334) being particularly important on both substrates (fitness ≤ −1.8 in all CS and HA experiments) (Figure 6A). Other components of PUL57 were preferentially important on one substrate only. BT3349 (a sulfatase) was only important on CS, as expected given that CS contains sulfated sugars, which are not present in HA. The recently described polysaccharide lyase family PL29 enzyme encoded by BT3328 was only important for fitness on HA, despite the activity of this enzyme against both HA and CS *in vitro* ^49^.

**Figure 6.**
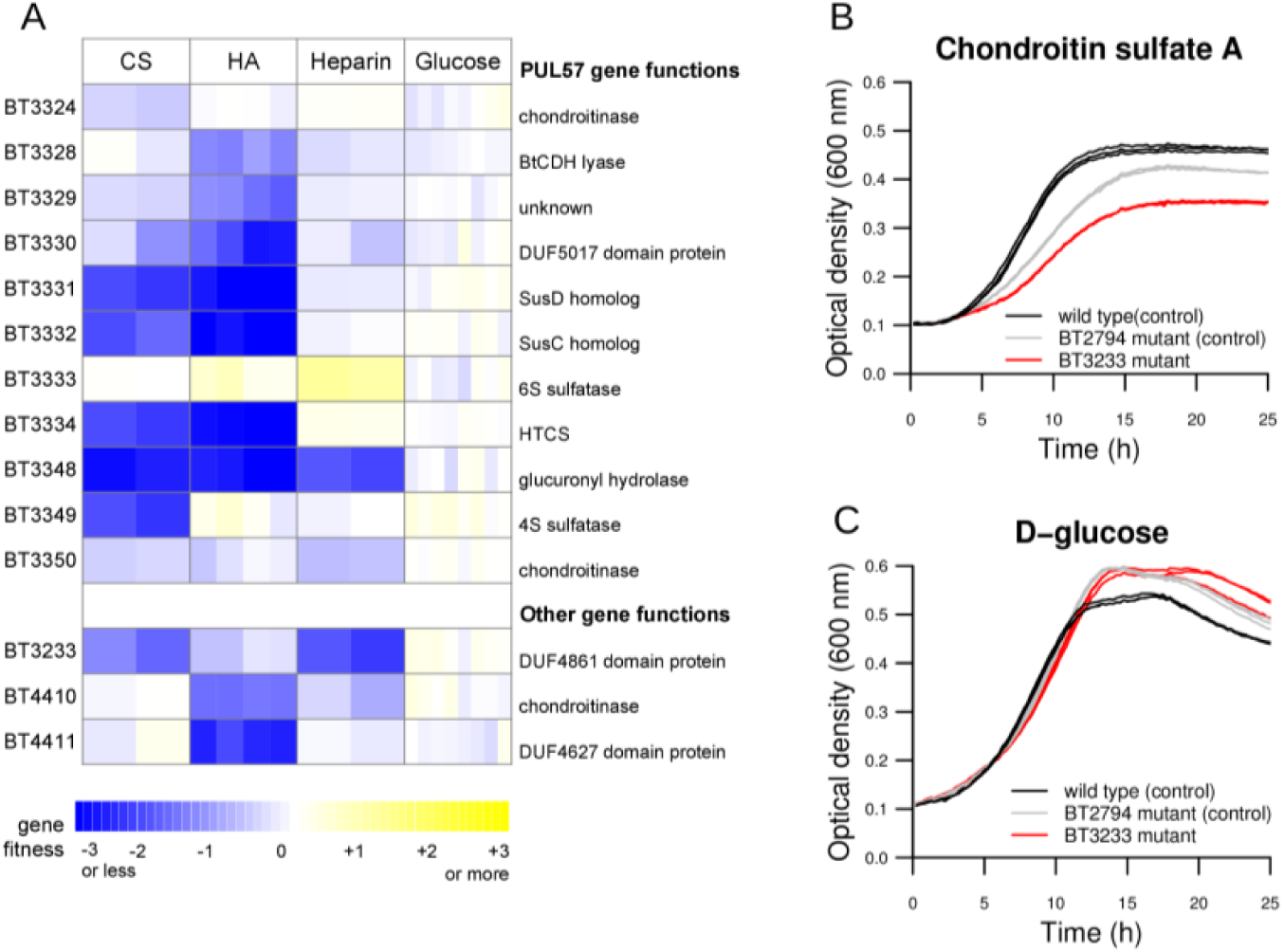
BT3233 and BT4410 are important for glycosaminoglycan utilization. (A) Mutant fitness data for the utilization of chondroitin sulfate A (CS), hyaluronic acid (HA), heparin, and glucose. Replicate experiments are displayed separated in the heatmap. (B,C) Growth curves of wild-type *B. thetaiotaomicron*, a BT3233 transposon mutant, and a BT2794 transposon mutant in (B) CS and (C) glucose (in which no phenotypes were predicted). The BT2794 mutant was used as a control, as this gene does not have a phenotype during growth on CS, unlike BT3233. The reduced growth of the BT2794 control relative to wild-type likely reflects the burden of expressing the erythromycin antibiotic resistance gene during growth in CS.

The first new GAG utilization gene we identified was BT4411, which was important for growth on HA but not on CS (Figure 6A). BT4411 is predicted to be in the same operon as the CS lyase BT4410, which was also more important for fitness on HA. BT4411 contains a domain of unknown function, DUF4627, which to our knowledge has not been previously characterized. DUF4627 has significant similarity to CBM_4_9 (carbohydrate binding domain PF02018) ^50^, the BT4410-4411 operon is conserved in several other *Bacteroides* species, and both proteins have putative signal peptides ^51^. Based on these data, we speculate that BT4410 and BT4411 form a complex and that BT4411 contributes to substrate binding.

Finally, we found that BT3233 was important for growth on 3 GAGs: CS (fitness = −1.8 and −1.4 in two experiments), HA (fitness = −0.7 in two experiments), and heparin (fitness = −2.3 and −2.0 in two experiments) (Figure 6A). BT3233 is a member of DUF4861, an uncharacterized protein family with predicted carbohydrate binding activity ^50^. Homologs of BT3233 are mostly confined to *Bacteroides* species, including a number of human commensal species. Using an individual transposon mutant from our archived collection, we confirmed that BT3233 is important for growth on CS (Figure 6B,C). Although its precise function is unknown, our data suggests that BT3233 plays an important role in the catabolism of diverse GAGs. Overall, our investigation of novel polysaccharide utilization genes identified three new genes with roles in GAG catabolism, including two genes with domains of unknown function.

### Disaccharide catabolism through a putative 3-keto intermediate

We observed that components of a gene cluster (BT2157, BT2159, and BT2160) were important for catabolizing the disaccharides trehalose (glucose-α-1,1-glucose), leucrose (glucose-α-1,5-fructose), palatinose (glucose-α-1,6-fructose), and the trisaccharide raffinose (galactose-α-1,6-glucose-β-1,2-fructose) (Figure 7A). BT2160 encodes a SusR-family transcription factor that likely regulates the cluster. In support of this view, a previous study used comparative genomics to identify a binding motif for BT2160, and predicted that this transcription factor only regulates target genes within its own predicted operon (BT2156-2160) ^13^, although a biological role for this system was not identified. BT2156 did not have a significant phenotype under any of our assayed conditions. This lack of phenotypes was potentially due to genetic redundancy, as *B. thetaiotaomicron* contains a second gene cluster (BT4446-4449) that is highly similar to BT2156-2159 (Supplementary Figure 1), except that BT4446-4449 does not contain a SusR-family regulator nor does it have a significant phenotype in our dataset. BT2158 is very similar to BT4448 (the nucleotide sequences are 98% identical), and we were not able to unambiguously map transposon insertions within these genes using our standard mapping parameters. To determine if BT2158 was important for fitness, we re-ran our transposon insertion mapping with less stringency by only requiring one nucleotide difference to assign insertions to BT2158 and BT4448, which successfully generated fitness data for both of these genes. While mutants of BT4448 did not have strong phenotypes, BT2158 was important for growth on trehalose, leucrose, palatinose, and raffinose (fitness ≤ −1.7 in all experiments on these substrates). Furthermore, BT2158 had the highest cofitness with BT2160 (*r* = 0.62). These results demonstrate that BT2157-2160 are all important for fitness on similar substrates, and thus are very likely to act in the same pathway.

**Figure 7.**
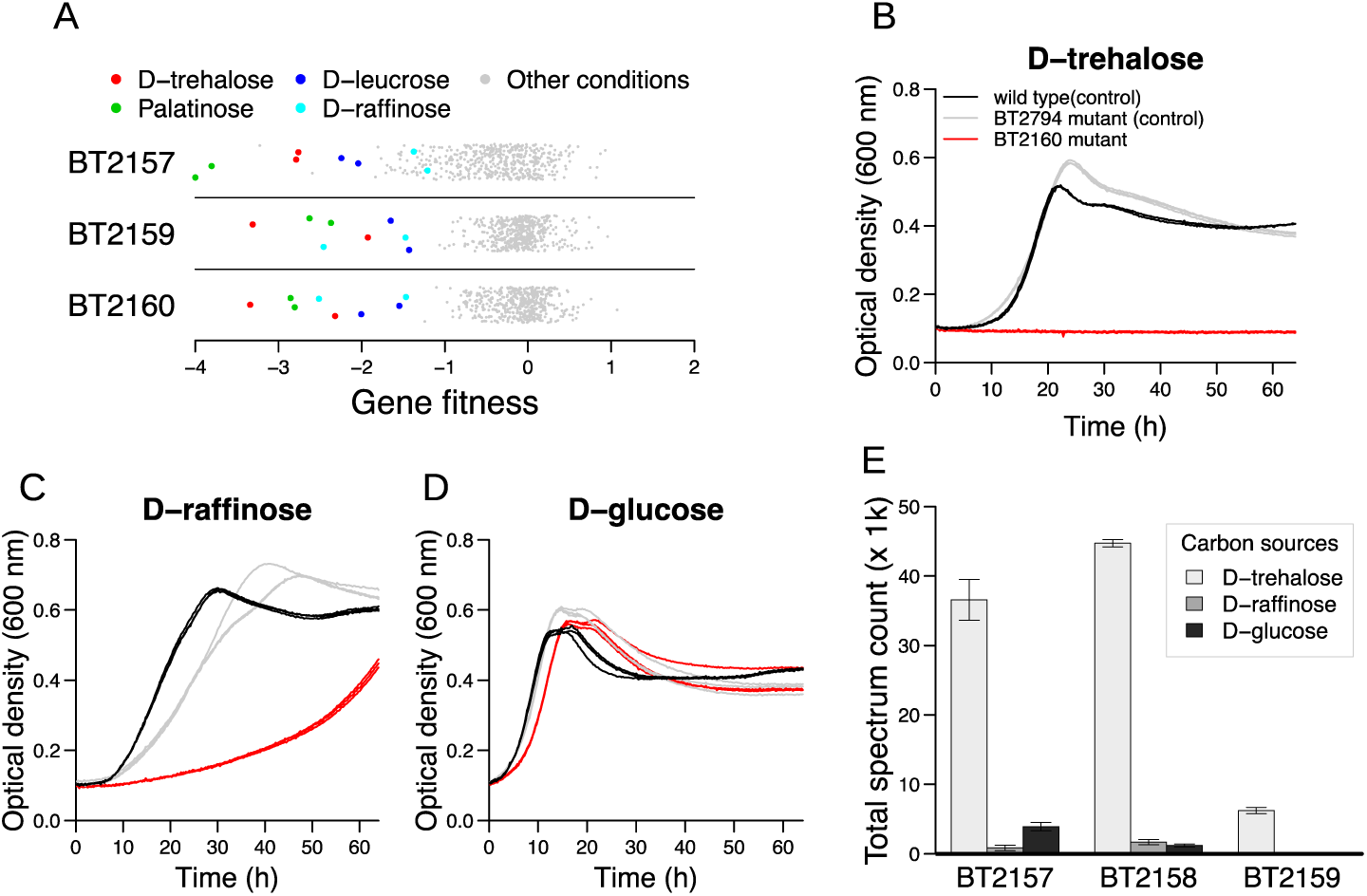
Identification of a gene cluster for di-and trisaccharide catabolism in *B. thetaiotaomicron.* (A) Gene fitness values for BT2157, BT2159, and BT2160 under all tested conditions. The *y*-axis is random. (B-D) Growth curves of wild-type *B. thetaiotaomicron*, a BT2160 transposon mutant, and a BT2794 transposon mutant in defined media with either trehalose (B), raffinose (C), or glucose (D) as the sole carbon source. BT2794 does not have a phenotype under any of these conditions in our pooled fitness experiments, and hence was used as a control. Each of three replicate growth curves are plotted (many replicates are highly overlapping). (E) Relative quantification of three proteins (BT2157-2159) that were predicted to be regulated by BT2160 after growth on the indicated carbon sources. Data are the average of 4 replicates. Error bars represent one standard deviation.

To further explore this system, we examined the growth of an individual BT2160 transposon mutant on different substrates. The mutant completely lost the ability to use trehalose as a carbon source, and had a substantially longer lag phase than the wild-type strain on raffinose (Figure 7B-D). One explanation for the mutant’s growth on raffinose is that the fructose monomer can be removed and consumed by the fructans PUL22, independent of the activity of BT2156-2160. Indeed, components of the fructans PUL are mildly important for consuming all 3 fructose-containing sugars (leucrose, palatinose, and raffinose; Figure 3 shows data for palatinose and raffinose), demonstrating that two different systems in *B. thetaiotaomicron* are required for optimal growth on certain small fructose-containing sugars. Next, we examined the induction of this operon on different substrates. A targeted proteomic analysis of stationary phase cultures revealed a significant increase in abundance for three of the target proteins (BT2157-2159) when grown on trehalose (Figure 7E). However, despite the importance of these genes for growth on raffinose, we did not observe increased abundance of these proteins in stationary phase of cultures grown on raffinose (Figure 7E).

Sequence analysis of the *B. thetaiotaomicron* BT2156-2159 proteins revealed similarities to some proteins putatively involved in the catabolism of disaccharides through a 3-ketoglycoside intermediate. Although we currently lack a complete understanding of the 3-ketoglycoside catabolic pathway, biochemical evidence for the oxidation of disaccharides by *Agrobacterium tumefaciens* has long been known ^52^. Furthermore, a cloned genomic fragment of *A. tumefaciens* conferred sucrose-degrading capabilities to *E. coli*, and a number of biochemical activities were identified and used to propose a catabolic pathway: sucrose is first oxidized to 3-ketodisaccharide and hydrolyzed in the periplasm, and then the 3-ketoglucose is transported into the cytoplasm and reduced (Supplementary Figure 2) ^53^. Although this genomic fragment was not sequenced, it did reveal that these biochemical activities are encoded within a single gene cluster in some bacteria. Similar evidence for disaccharide catabolic pathways via a 3-ketoglycoside intermediate have been observed for trehalose utilization in *Sinorhizobium meliloti* ^54^ and lactose utilization in *Caulobacter crescentus* ^55^, with the latter example expected to involve a 3-ketogalactose intermediate. Only the first step of this proposed pathway has been definitively linked to genes. This disaccharide 3-dehydrogenase is encoded by two unrelated gene clusters, *thuAB* from *S. meliloti* and *lacABC* from *C. crescentus*. The 3-ketoglycoside hydrolase and the 3-ketosugar reductase activities have not, to our knowledge, been associated to genes in any bacterium.

Here, we propose that *B. thetaiotaomicron* catabolizes disaccharides through a 3-ketoglycoside intermediate, and that BT2156-2160 are involved in this process. Our evidence relies on two observations. First, by mining fitness data from a previous large survey ^7^, we identified shared genes that are chromosomally clustered with *lacABC* and are important for catabolizing diverse glycosides in *C. crescentus, Pedobacter* sp. GW460-11-11-14-LB5, and *Echinicola vietnamensis* KMM 6221. In addition to the known LacABC disaccharide dehydrogenase, we found that one or more IolE-like sugar epimerases, MviM-like dehydrogenases, MFS transporters, and DUF1080-containing proteins were important for fitness on different disaccharides, raffinose, and/or salicin (a glucoside of glucose and salicyl alcohol) in these three species (Supplementary Figure 3). In addition, in each of these three species, at least one copy of all of these proteins is encoded within a single gene cluster, although in some instances a paralogous protein outside the cluster has a stronger or variable phenotype (Supplementary Figure 1).

Our second line of evidence is that proteins in BT2156-2160 are similar to proteins from the *C. crescentus, Pedobacter* sp. GW460-11-11-14-LB5, and *E. vietnamensis lacABC* gene clusters, even though *B. thetaiotaomicron* does not contain the full gene cluster (Supplementary Figure 1). BT2156 encodes a predicted IolE-like epimerase, BT2157 a DUF1080-containing protein, and BT2158 and BT2159 encode MviM-like dehydrogenases. BT2158 is likely localized to the periplasm ^51^, while BT2159 is expected to be in the cytoplasm. According to Pfam ^50^, DUF1080 is a hydrolase with structural similarity to endo-1,3-1,4-β-glucanase. Given the importance of various members of this protein family for consuming different glucosides in diverse bacteria, and the genomic proximity of this protein family with other components of the 3-ketoglycoside pathway, we propose that DUF1080-containing proteins are more specifically 3-ketoglycoside hydrolases. In addition, DUF1080-containing proteins are likely localized to the periplasm, as they often contain a putative signal peptide ^51^. While the exact details remain to be elucidated, we propose that in *C. crescentus, Pedobacter* sp. GW460-11-11-14-LB5, and *E. vietnamensis*, various glucosides are oxidized by LacABC to a 3-ketoglucoside in the periplasm, hydrolyzed to a 3-ketosugar and a second monomer by a DUF1080 hydrolase in the periplasm, and the 3-ketosugar is taken up via the MFS transporter and further metabolized by the IolE-like epimerase and MviM-like dehydrogenase. In *B. thetaiotaomicron*, we have no genetic evidence for the 3-glucoside dehydrogenase nor the MFS transporter, so it remains unclear how the 3-ketoglucoside is formed and where this metabolism occurs.

### An efflux system for bile salts

During growth in the human gut, *B. thetaiotaomicron* is exposed to complex polysaccharides and also to bile salts, which can inhibit bacterial growth. Bile acids are acidic steroids that are synthesized by the liver from cholesterol. Primary bile acids are conjugated with taurine or glycine before secretion, and the conjugation step greatly increases their water solubility under physiological pH, thus the conjugated form of bile acids are usually called bile salts. Once they reach the colon, bile salts can be deconjugated and oxidized by bacteria into so-called secondary bile acids. Bile salts play an important role as an innate immune defense through their potent antibacterial activity, primarily by disrupting bacterial membrane integrity ^56,57^. Consequently, commensal bacteria have evolved a number of strategies to cope with bile salts including removal by efflux pumps, deconjugation by bile salt hydrolases (BSH) and oxidation and epimerization of hydroxyl groups by hydroxysteroid dehydrogenases (HSDH) ^57^. BSH and HSDH homologs have been identified in *B. thetaiotaomicron* ^58^, and BT2086 has recently been demonstrated to have BSH activity *in vitro* and *in vivo* ^59^.

To identify *B. thetaiotaomicron* genes involved in bile salt tolerance, we examined fitness data from growing the mutant library in the presence of inhibitory concentrations of either the primary bile acid chenodeoxycholic acid, the secondary bile acids lithocholic acid or deoxycholic acid, or cholic acid-derived bile salts (a mixture of sodium cholate and sodium deoxycholate, which we refer to here as “bile salts”). Furthermore, we performed these experiments in two different growth media: BHIS and VB. From these data, we identified a predicted four-gene operon BT2792-2795 that was important for fitness under deoxycholic acid, chenodeoxycholic acid, and bile salt stress (Figure 8A). These genes were important for fitness in both growth media and at multiple stressor concentrations; the phenotypes were more pronounced at higher concentrations. For example, at 0.5 mg/mL bile salts, the fitness values for all four genes ranged from −2.1 to −3.3 (irrespective of growth media). These genes were not important when stressed with lithocholic acid, at least under the concentrations we assayed. Based on homology ^60^, these proteins form a tripartite multidrug efflux system, with BT2793 as the inner membrane component, BT2794 as the membrane fusion component, and BT2795 as the outer membrane component. BT2792 is a transcriptional regulator that likely regulates the operon. Interestingly, these genes did not have a significant phenotype in any of the other conditions that we profiled (Figure 8A), including dozens of diverse antibiotics, suggesting that this efflux system is specific to bile acids and bile salts. Closely related efflux systems are present in other *Bacteroides* species ^61^, although the importance of those genes for bile salt tolerance remains to be determined.

**Figure 8.**
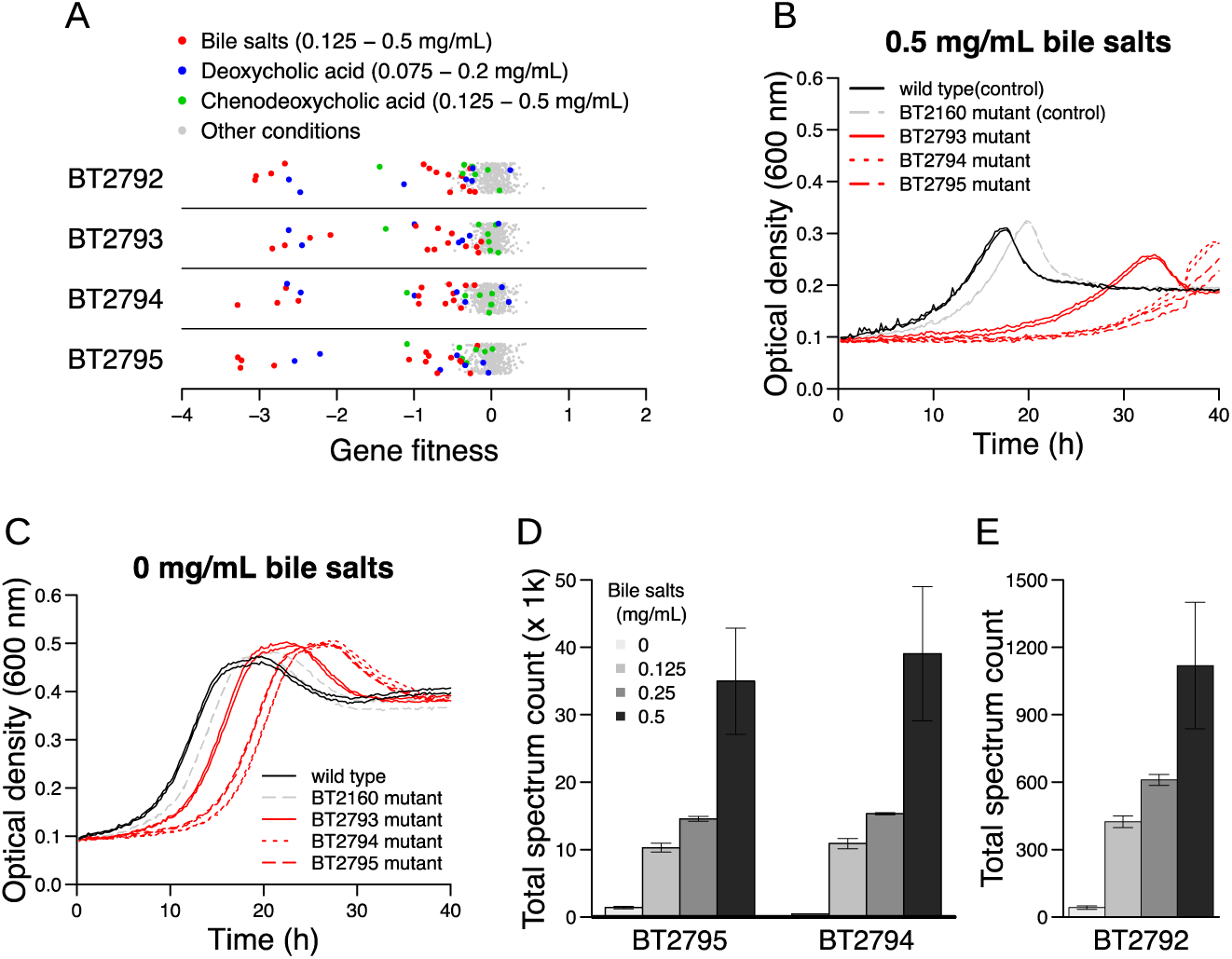
Identification of an efflux system for bile salts in *B. thetaiotaomicron*. (A) Gene fitness values for BT2792-2795 across all experiments with bile salts and bile acids conditions highlighted. The highlighted experiments without a significant phenotype correspond to lower concentrations of the compound. The *y*-axis is random. (B,C) Growth curves of individual *B. thetaiotaomicron* transposon mutant strains in VB medium with (B) 0.5 mg/mL bile salts or (C) no added stress. The BT2793-2796 mutants all have extended lag times compared with controls. For each strain/growth condition, we show two replicate growth curves. (D,E) Relative abundance of proteins (D) BT2794 and BT2795 or (E) BT2792 from wild-type cells grown in BHIS with varying concentrations of bile salts demonstrate their upregulation by bile salts in a concentration-dependent manner. Data are averages of four replicate experiments and error bars represent one standard deviation.

To confirm the contribution of this operon to bile salt tolerance in *B. thetaiotaomicron*, we assayed the growth of individual transposon mutants from our arrayed collection. These data revealed that mutants of BT2793, BT2794, and BT2795 all had longer lag phases and reduced growth rates relative to control strains when stressed with 0.5 mg/mL bile salts (Figure 8B,C). To determine if this efflux system is induced by the stressor, we performed targeted proteomic analysis using wild-type cells grown in the presence of various concentrations of bile salts. BT2792, BT2794, and BT2795 were all induced in the presence of bile salts, and the amount of induction increased with higher bile salt concentrations (Figure 8D). Bile salt-mediated upregulation of efflux pumps has also been observed in other bacteria from the human microbiota ^62-65^. Taken together, our data shows that the efflux system encoded by BT2792-2795 is regulated by and important for resistance to bile salts, suggesting that this efflux system in *B. thetaiotaomicron* has evolved to specifically mitigate bile salt stress in the human intestinal tract.

### Transport of antibiotics and biocides

Given the extensive use of antimicrobial compounds to treat infection, it is expected that commensal members of the human microbiota are exposed to these inhibitors. Indeed, antibiotic treatment has been shown in some instances to decrease the relative abundance of *Bacteroides* species ^66^. We examined the fitness data generated from assaying the mutant library during growth in the presence of different classes of inhibitory compounds including antibiotics and biocides. Given the prevalence of compound efflux as a mechanism of antibiotic resistance in bacteria ^67^, we focused our analysis on transport-related proteins with strong phenotypes on different inhibitors. For example, BexA (BT0588) belongs to the multidrug and toxic compound extrusion (MATE) family and has previously been shown to be responsible for the efflux of the fluoroquinolone antibiotics norfloxacin and ciprofloxacin in *B. thetaiotaomicron*^68^. In our dataset, we found a mild fitness defect for BT0588 mutants in the presence of ciprofloxacin (minimum fitness on ciprofloxacin = −1.6, although it was not statistically significant). The loss of BT0588 resulted in a more pronounced phenotype with another fluoroquinolone, lomefloxacin, or with paraquat (minimum fitness in a lomefloxacin experiment = −3.1; minimum fitness in a paraquat experiment = −3.6) (Figure 9A). These results illustrate the utility of fitness profiling across many inhibitory compounds to associate new antibiotics to a previously studied efflux protein.

**Figure 9.**
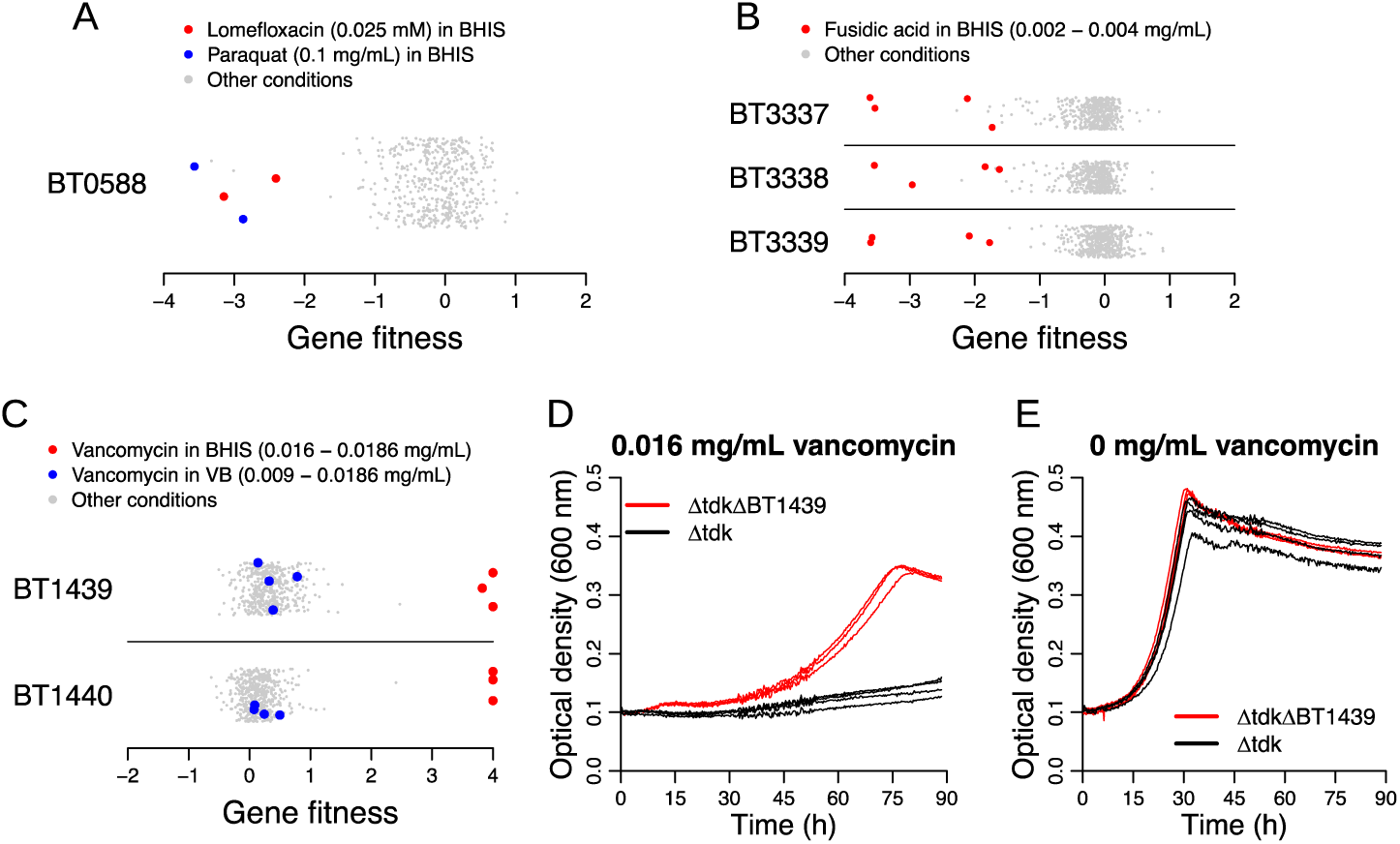
Identification of antibiotic resistance genes in *B. thetaiotaomicron.* (A-C) Gene fitness values for BT0588 (also known as BexA) (A), BT3337-3339 (B), and BT1439-1440 (C) under all tested conditions. The *y*-axis in these plots is random. (D,E) Growth curves of a BT1439 deletion and the parental strain (Δ*tdk*) in the presence (D) or absence (E) of vancomycin in BHIS medium. The BT1439 deletion strain exhibits more growth than the parental strain in the presence of vancomycin. Three replicate growth curves per strain/condition are displayed.

We also identified phenotypes for previously unstudied transport systems. For example, we found that the RND transporter system encoded by BT3337-3339 was important for fusidic acid tolerance (Figure 9B), and to a lesser extent for tolerating other inhibitors, including cefoxitin and the antipsychotic drug chlorpromazine. These results suggest that the substrate range of BT3337-3339 is broad and includes antimicrobials of diverse chemical classes.

Lastly, we identified a predicted two-gene operon BT1439-1440 with high positive fitness values when grown in the presence of vancomycin (Figure 9C), a glycopeptide antibiotic that inhibits the biosynthesis of peptidoglycan in the periplasmic space. Vancomycin is commonly used for treating *Clostridium difficile* infections (CDI), and often taken orally. One study showed that fecal levels of vancomycin in CDI patients can range from 0.05 mg/mL to 2 mg/mL during the antibiotic treatment ^69^, which is above the inhibitory concentration we used in our fitness assays (0.009-0.0186 mg/mL). These numbers suggest that vancomycin use in humans can impact *B. thetaiotaomicron* in the gut. BT1439 is a SusD homolog and BT1440 is a SusC homolog, and these genes have high cofitness (*r* = 0.76), indicating that these proteins likely act together. Although SusCD complexes are best known to be involved in translocating carbohydrate substrates from outside the cell to the periplasmic space, no growth deficiency was found in any of the carbohydrate conditions that we assayed. To validate our genome-wide data, we assayed the growth of a BT1439 deletion mutant and found that it was substantially more resistant to vancomycin relative to the parental control (Figure 9DE). The concentration of vancomycin that inhibited growth by half (IC_50_) was over seven times higher in the BT1439 mutant compared to the non-mutated control (0.0047 versus 0.00066 mg/mL) (Supplementary Figure 4). Interestingly, the vancomycin phenotype of BT1439-1440 was only observed in BHIS media, and not in VB media (Figure 9C). We propose that this media-specific difference in phenotype could be due to altered regulation or the impact of vancomycin binding to another compound in the growth media. Regardless, our data reveal surprising links between genes annotated as carbohydrate transporters and antibiotic sensitivity.

## Discussion

A small fraction of the gene content of the human microbiome has been experimentally characterized, and as such, there are large knowledge gaps in our molecular understanding of these bacteria. Here, we address this knowledge gap using high-throughput genetics in *B. thetaiotaomicron*, the most commonly studied gut commensal bacterium. We performed hundreds of chemical-genetic experiments with a randomly barcoded transposon mutant library and generated nearly two million gene-phenotype measurements. Using these data, we identified novel phenotypes for PULs, linked proteins with domains of unknown function to polysaccharide and disaccharide degradation, and identified an efflux system specific for bile salts tolerance. This large dataset contains phenotypes for hundreds of additional *B. thetaiotaomicron* genes and should be an invaluable resource for gene function inference and hypothesis generation for the scientific community. In particular, the 612 genes with either a specific phenotype or high cofitness to another gene are attractive candidates for follow-up investigations. To facilitate these future studies, the *B. thetaiotaomicron* fitness data is available for comparative analyses at the Fitness Brower (http://fit.genomics.lbl.gov), a web portal with data from over 5,000 genome-wide fitness experiments from 35 bacteria.

Much of the research to date on *B. thetaiotaomicron* has focused on its ability to degrade complex polysaccharides using PULs. While we were able to link 20 PULs to carbon sources via specific phenotypes, this number represents less than 25% of the total number of predicted PULs in the genome. There are a number of potential reasons for our inability to detect a carbon source phenotype for many PULs. First, *B. thetaiotaomicron* may contain two or more PULs that act on the same substrate, and this genetic redundancy will mask the impact of any single-gene loss-of-function mutations. To address this redundancy, future efforts could investigate genetic interactions via double mutants, or by interrogating carbon source utilization in other *Bacteroides* strains under the expectation that not all of these strains will have redundant systems. Indeed, it is known that different gut *Bacteroides* species have different polysaccharide-degrading capabilities ^46^. Second, the PUL (or individual genes within the PUL) may perform its activity extracellularly, such that other mutants in the pooled library can complement any putative growth deficiency *in trans*. In addition to assaying mutants individually (including those in our archived mutant collection), newer approaches for performing genome-wide fitness assays within droplets are becoming available, which hold the potential for eliminating complementation by other mutant strains within the pooled library ^70^. Third, some of the PULs may not be expressed in our simple growth conditions or are involved in the breakdown of polysaccharides that we did not profile. Fourth, some of these PULs may not have evolved to eat complex polysaccharides, but rather simpler sugars and oligosaccharides. For example, we identified a PUL (BT3567-3569) that was only important for fitness on the disaccharide laminaribiose, suggesting that chemical-genetic profiling with additional simple carbon sources may reveal new functions for some of these PULs. Lastly, the definition of a PUL is very minimal, and only requires a SusCD gene pair. Some of these minimal “PULs” may have other cellular roles. For instance, we showed that a two-gene PUL (BT1439-1440) confers susceptibility to vancomycin in BHIS, while we were unable to detect a carbon source phenotype for this system.

Our genetic approach can be applied generally to discover new gene functions in the human microbiome. In particular, the barcoding of mutant libraries enables the rapid assessment of gene importance across many conditions at low cost ^7^. Nevertheless, challenges still need to be overcome, including the accelerated development of genetic tools for non-model species of the human microbiota ^71,72^, and the development of scalable phenotypic assays that more accurately reflect the natural ecology of these bacteria, including interactions with other bacteria, phage, and the human host. In addition, the conservation of a phenotype in multiple bacteria is a powerful indicator of gene function ^7^, and the generation of similar gene-phenotype maps in related bacteria will expand the scope of analysis. In particular, extending our approach to additional *Bacteroides* species from the human gut will provide higher confidence inferences of gene function for conserved genes, as well as provide insights into the functions of accessory genes that are present in a subset of the genomes. A greatly improved understanding of the genes in the human microbiota will accelerate the development of interventions and therapeutics to improve human health.

## Materials and Methods

### Strains, plasmids, and compounds

The strains, plasmids, and oligonucleotides used in this study are listed in Supplementary Tables 5-7. All oligonucleotides and gBlocks were ordered from Integrated DNA Technologies. All PCR reactions were carried out with Q5 hot start DNA polymerase from New England Biolabs (NEB). PCR products were cleaned with the DNA Clean & Concentrator kit (Zymo Research). Gibson assembly reactions were performed with the Gibson Assembly Master Mix from NEB. Golden Gate assembly enzymes Esp3I (BsmBI) and BpiI (BbsI) were purchased from ThermoFisher Scientific. T4 DNA ligase and buffer were purchased from NEB. Plasmid isolation was accomplished using the QIAprep Spin Miniprep Kit (Qiagen). In this study, we used many of the compounds that we previously reported in an investigation of bacterial mutant fitness ^7^. We also purchased and assayed additional compounds that are specifically relevant to *B. thetaiotaomicron*, such as polysaccharides and bile salts. Given that some of the polysaccharides contain heterogeneous compounds, we provide detailed vendor information in Supplementary Table 4.

### Constructing a barcoded transposon delivery vector for *B. thetaiotaomicron*

We previously described a *mariner* transposon delivery vector library pHLL255 (magic pool) containing hundreds of combinations of promoters and erythromycin drug markers ^26^. The plasmids in the magic pool contain DNA barcodes that enable the parallel tracking of mutagenesis efficiency for each plasmid design in a single assay. To identify a suitable vector for mutagenizing *B. thetaiotaomicron*, we first constructed a small mutant library by conjugating with an *E. coli* strain carrying pHLL255. We performed TnSeq on this library to map transposon insertion locations and their associated DNA barcodes, enabling us to infer which vector design in the magic pool resulted in the mutation ^26^. Analysis of this library revealed that the optimal drug marker for mutagenizing *B. thetaiotaomicron* was the erythromycin resistance gene *ermG* ^20,73^, the optimal promoter for expressing the *mariner* transposase was the native *rpoD* (BT1311) promoter ^20^, and the optimal promoter for expressing *ermG* was the native *dnaE* (BT2230) promoter.

Although this vector design had the highest insertion efficiency relative to the other designs in the magic pool, it still resulted in a significant strand bias whereby the insertion orientation of the transposon was more often in the same transcriptional orientation as the mutated gene. Based on our previous work, we proposed that stronger expression of the selection marker on the transposon might eliminate this strand bias ^26^. So, in addition to constructing the best but potentially suboptimal vector design from the magic pool results (pTGG45), we also constructed another vector (pTGG46) by replacing the *dnaE* promoter in pTGG45 with a *cepA* hybrid promoter from *Bacteroides fragilis* RBF-103. This hybrid promoter contains a putative insertion sequence element (IS1224), and has been shown to have increased promoter activity relative to the native *cepA* promoter ^74^. To determine which vector was the best combination of high efficiency and low strand bias, we constructed two small mutant libraries with either pTGG45 or pTGG46. TnSeq analysis of these libraries revealed that both had high insertion efficiency, but while the strand bias was again significant for pTGG45 (60% of insertions were in the same orientation as transcription of the mutated gene), there was essentially no strand bias for pTGG46 (49.4%). We then barcoded pTGG46 (pTGG46_NN1) with millions of random 20-nucleotide barcodes using Golden Gate assembly, as previously described ^26^.

### Transposon mutant library construction

We carried out transposon mutagenesis by conjugating *B. thetaiotaomicron* recipient cells with *E. coli* donor cells carrying the barcoded transposon vector library pTGG46_NN1 at 1:1 ratio. Specifically, we grew 10 mL of wild-type *B. thetaiotaomicron* in BHIS overnight at 37 °C. The next morning, we recovered a 2 mL freezer stock of AMD776 (*E. coli* WM3064 with pTGG46_NN1) in 50 mL LB supplemented with 50 μg/mL carbenicillin and 300 μM diaminopimelic acid (DAP) at 37 °C. When the OD_600_ of the *E. coli* donor strain reached ~1.0, we harvested 15 OD_600_ units of the culture and washed the cells 3 times with fresh BHIS supplemented with DAP. Then, 15 OD_600_ units of *B. thetaiotaomicron* wild-type cells were harvested and mixed with the washed donor strain cells, and resuspended in a final volume of 60 μL with BHIS supplemented with DAP. The resuspension was spotted onto 0.45-µm gravimetric analysis membrane filters (Millipore), and incubated aerobically overnight on BHIS agar plates supplemented with DAP at 37 °C. The next day, the conjugation mixture was scraped from the membrane and resuspended into 6 mL fresh BHIS medium supplemented with 10 μg/mL erythromycin, and plated at different dilutions on BHIS plates supplemented with 10 μg/mL erythromycin. The plates were incubated at 37 °C for 48 hours to let visible colonies develop. We then pooled ~250,000 colonies, and grew the pooled library in liquid BHIS supplemented with 10 μg/mL erythromycin and 200 μg/mL gentamicin (to further select against the *E. coli* conjugation donor strain). When the cells reached saturation, we made multiple, single-use glycerol stocks of the final library and extracted genomic DNA for TnSeq analysis. To map the genomic location of the transposon insertions and to link these insertions to their associated DNA barcode, we used the same TnSeq protocol that we described previously ^25^. The final mutant library was named Btheta_ML6. Lastly, using single-cell sorting, we generated and mapped an archived collection of individual transposon insertion mutants from this library (A.L.S, A.M.D, and K.C.H, in review).

### Individual mutant strain construction and assays

We constructed an unmarked, in-frame deletion mutant of BT1439 using a previously established approach ^75^. The individual transposon insertion mutants we assayed were from the arrayed collection. We performed growth curves of individual mutants in a 96 well plate in a Tecan Infinite F200 instrument housed in a Coy anaerobic chamber. To calculate IC_50_ values for the BT1439 deletion mutant and parent strain on vancomycin, we used the drc package in R ^76^. Specifically, we used drm() with the LL.4() model and defined the response as the average OD_600_ up until the time that uninhibited cells (no vancomycin) reached their maximum OD_600_ value.

### Growth conditions

Growth media was purchased from BD and most compounds were purchased from Sigma-Aldrich. Some carbohydrates were purchased from other vendors (Supplementary Table 4). We typically grew *B. thetaiotaomicron* anaerobically in a Coy chamber at 37 °C with one of two different growth media. First, we used Brain Heart Infusion Supplemented (BHIS) medium using Bacto Brain Heart Infusion (BHI) as the base. Second, we used a defined medium developed by Varel and Bryant (VB) ^77^. The recipes for both media are provided in Supplementary Table 8. For culturing individual *B. thetaiotaomicron* transposon insertion mutants, we supplemented the media with 10 μg/ml erythromycin. For growth on solid media, we used 1.5% (w/v) agar.

### Genome-wide mutant fitness assays

We performed genome-wide mutant fitness assays as described previously ^7^. Briefly, we thawed an aliquot of the full transposon mutant library, inoculated the entire aliquot into 50 mL of BHIS supplemented with 10 μg/ml erythromycin, and grew the library anaerobically at 37°C until the cells reached mid-log phase. We then collected 6 cell pellets of ~1.0 OD_600_ unit each (the “Time0” samples; we collect multiple pellets for the Time0 sample because we perform BarSeq sequencing on each of these). We then used the remaining cells to inoculate competitive growth assays. Nearly all fitness assays were performed in 1.2 mL of growth medium in either a 24-well transparent microplate (Greiner) or in a 96-deepwell plate (Costar). For fitness assays in BHIS or VB, we diluted the recovered mutant library to a starting OD_600_ of 0.02 for each experiment. For carbon source assays in VB, we washed the cells in VB lacking a carbon source two times. For nitrogen source experiments, we washed the cells in VB lacking a nitrogen source two times. For all growth-based fitness assays, we grew the cells until the cultures reached stationary phase, and then collected cell pellets (the “Condition” sample). We also performed a few “survival” experiments in which we incubated the mutant library aerobically for different amounts of time. After this incubation, we inoculated cells into fresh BHIS and grew anaerobically to select for cells that survived the stress. We extracted genomic DNA from the Time0 and Condition samples using either the DNeasy Blood and Tissue kit (Qiagen) or in a 96-well microplate format with the QIAamp 96 DNA Qiacube HT kit (Qiagen).

We performed barcode sequencing (BarSeq) as previously described ^7,78^. For most experiments, we performed BarSeq with indexed P2 oligos and a mix of non-indexed P1 oligos with variable lengths of N’s (2-5) to stagger the reads. For some experiments, we used BarSeq oligos with both P1 and P2 indexed to minimize the impact of incorrectly assigned indexes in Illumina HiSeq4000 runs ^79^. Strain and gene fitness scores were calculated as previously described ^25^. Briefly, strain fitness was calculated as the log_2_ ratio of barcode abundance after growth selection (Condition) versus before growth selection (Time0). Gene fitness was calculated as the weighted average of individual strain fitness values and was normalized to remove the effect of chromosomal position (due to variation in copy number along the chromosome). Finally, gene fitness was normalized so that the mode of the distribution was 0.

### Analysis of fitness data

Overall, we attempted 772 genome-wide fitness assays in this study, of which 492 passed our metrics for internal consistency ^25^. From these 492 experiments, we identified genes with a specific phenotype in only one or a handful of conditions using previously described criteria ^7^: |fitness| > 1 and |*t*| > 5 and 95% of |fitness| < 1 and |fitness| > 95th percentile(|fitness|) + 0.5, where *t* is fitness/estimated standard error and |*t*| > 5 ensures that the phenotype is statistically significant. For the analysis of the total number of genes with a specific phenotype, we excluded one gene that had specific phenotypes in control experiments with plain BHIS medium. A specific important phenotype has the same criteria as above except that the fitness score has to be negative. We identified pairs of genes with highly correlated patterns of phenotypes across all 492 successful experiments (cofitness) using the linear (Pearson) correlation and a threshold of 0.8 ^7^.

COG assignments and functional codes were downloaded from MicrobesOnline ^61^. Polysaccharide utilization loci assignments were derived from the CAZy database ^39^. Most data analyses to infer the putative functions of genes were performed with the Fitness Browser (http://fit.genomics.lbl.gov).

### Targeted proteomics experiments

For carbon source experiments, we grew wild-type *B. thetaiotaomicron* in VB with 20 mM D-trehalose, D-raffinose, or D-glucose. After overnight incubation, the OD_600_ values of the cultures were between 1.0 and 1.5, and the cultures were used for proteomics analysis. For bile salt experiments, we grew wild-type *B. thetaiotaomicron* in defined VB medium with 20 mM D-glucose and supplemented with bile salts at 4 different concentrations: 0, 0.125, 0.25, and 0.5 mg/mL. These cultures were growth for 20 hours, after which the OD_600_ values were between 0.5 and 0.9. For all experiments, we harvested 2 OD_600_ units of cells and washed two times with 10 mM NH_4_HCO_3_ before extracting the total protein using a methanol-chloroform precipitation method. Protein concentration was determined by Bio-Rad DC protein assay (Lot# 5000112) following manufacturer’s protocol, and the same amount of proteins were subjected to standard reduction, alkylation, and trypsin digestion procedures. A targeted selected reaction monitoring (SRM) method was developed to quantify unique peptides of proteins with the assistance of an in-house constructed *B. thetaiotaomicron* spectral library ^80^.

SRM targeted proteomic assays were performed on an Agilent 6460 QQQ mass spectrometer system coupled with an Agilent 1290 UHPLC system (Agilent Technologies, Santa Clara, CA). Twenty microgram peptides of each investigated sample were separated on an Ascentis Express Peptide C18 column (2.7-mm particle size, 160-Å pore size, 5-cm length × 2.1-mm inside diameter (ID), coupled to a 5-mm × 2.1-mm ID guard column with the same particle and pore size, operating at 60 °C; Sigma-Aldrich) operating at a flow rate of 0.4 mL/min via the following gradient: initial conditions were 98% solvent A (0.1% formic acid), 2% solvent B (99.9% acetonitrile, 0.1% formic acid). Solvent B was increased to 40% over 5 min, and then increased to 80% over 0.5 min, and held for 2 min at a flow rate of 0.6 mL/min, followed by a ramp back down to 2% solvent B over 0.5 min where it was held for 1 min to re-equilibrate the column to original conditions. The eluted peptides were ionized via an Agilent Jet Stream ESI source operating in positive-ion mode with the following source parameters: gas temperature = 250 °C, gas flow = 13 L/min, nebulizer pressure = 35 psi, sheath gas temperature = 250 °C, sheath gas flow = 11 L/min, capillary voltage = 3500 V, nozzle voltage = 0 V. Data were acquired using Agilent MassHunter v. B.08.02. Acquired SRM data were analyzed by Skyline v. 3.70 (MacCoss Lab Software).

### Strain, data, and code availability

The barcoded *B. thetaiotaomicron* transposon mutant library (Btheta_ML6) is available upon request. Fitness data from the 492 successful experiments is available for comparative analysis at the Fitness Browser (http://fit.genomics.lbl.gov). In addition, the fitness data (and all associated metadata) is available at: https://doi.org/10.6084/m9.figshare.7742855.v1. The software for RB-TnSeq is available at https://bitbucket.org/berkeleylab/feba.

## Acknowledgements

The *B. thetaiotaomicron* VPI 5482 *Δtdk* strain and the plasmid pExchange-tdk were kindly provided by the James A. Imlay lab at University of Illinois at Urbana-Champaign. We thank Surabhi Mishra for advice on constructing in-frame deletion mutants. This work was supported by Laboratory Directed Research and Development (LDRD) funding from Berkeley Laboratory, provided by the Director, Office of Science, of the US Department of Energy under contract DE-AC02-05CH11231, used resources of the Joint BioEnergy Institute supported by the Office of Science, Office of Biological and Environmental Research, of the US DOE under the same contract, and by the Allen Discovery Center at Stanford on Systems Modeling of Infection (to A.L.S. and K.C.H.). K.C.H. is a Chan Zuckerberg Investigator.

