## Supplemental Figures for "Large-scale chemical-genetics of the human gut bacterium *Bacteroides thetaiotaomicron*"

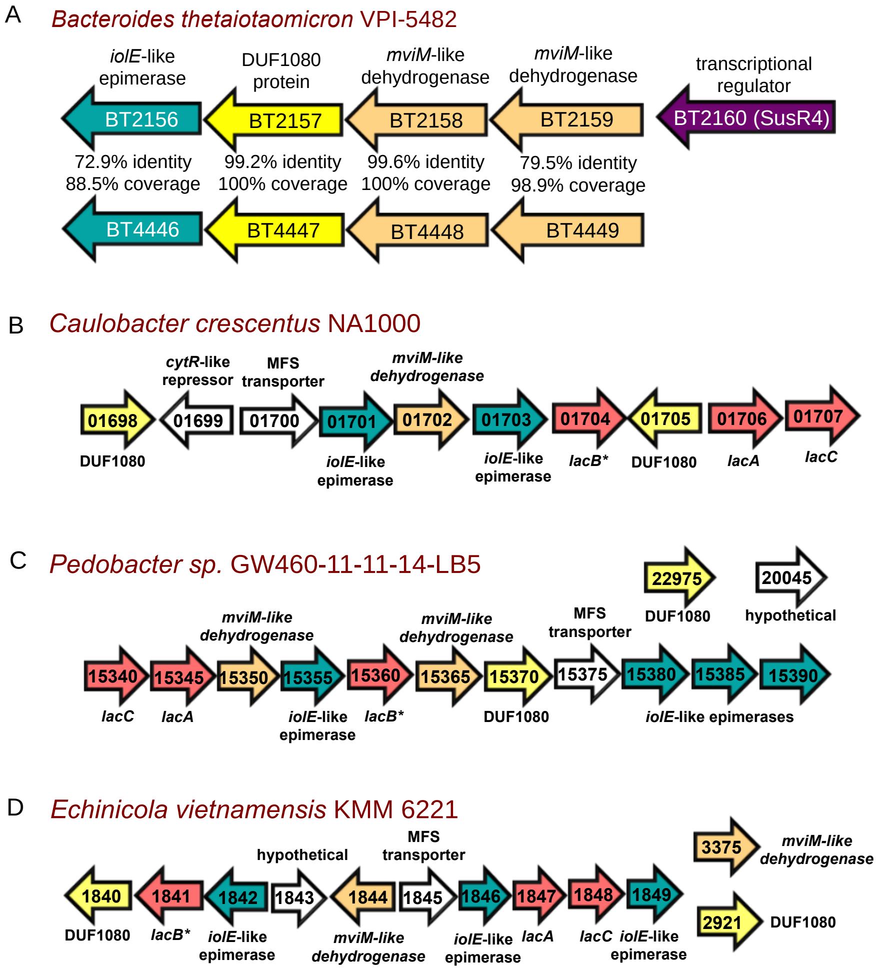


**Supplementary Figure 1. Organization of putative 3-ketoglycoside catabolism genes in four bacteria.** (A) *B. thetaiotaomicron*, (B) *Caulobacter crescentus*, (C) *Pedobacter* sp. GW460-11-11-14-LB5, (D) *Echinicola vietnamensis* KMM6221. Homologous genes between the four bacteria have the same color. The LacB proteins in *Pedobacter* sp. GW460-11-11-14-LB5 and *E. vietnamensis* KMM6221 are distantly related to characterized protein from *C. crescentus* (homology is not detectable by BLAST). We annotate them as LacB because they are also putative cytochrome c proteins (PFam PF00034) and are in the cluster. We do not show the prefixes for the genes: CCNA_ for *C. crescentus*, CA265_RS for *Pedobacter* sp. GW460-11-11-14-LB5, and Echvi_ for *E. vietnamensis* KMM6221. In (C) and (D), we show genes that are not in the cluster but have relevant mutant phenotypes. In (A), we also show a similar gene cluster in *B. thetaiotaomicron* (BT4446-4449) that is not important for disaccharide catabolism.

A Sucrose degradation pathway VII in *Agrobacterium tumefaciens*

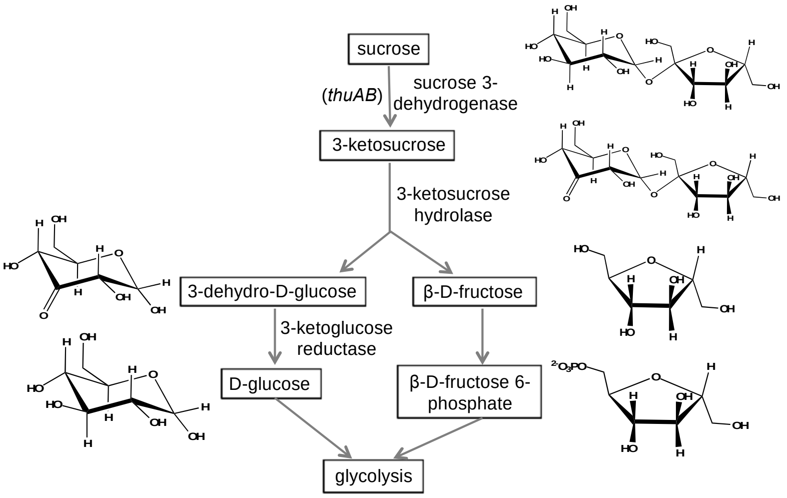


B Lactose degradation pathway II in *Caulobacter crescentus*

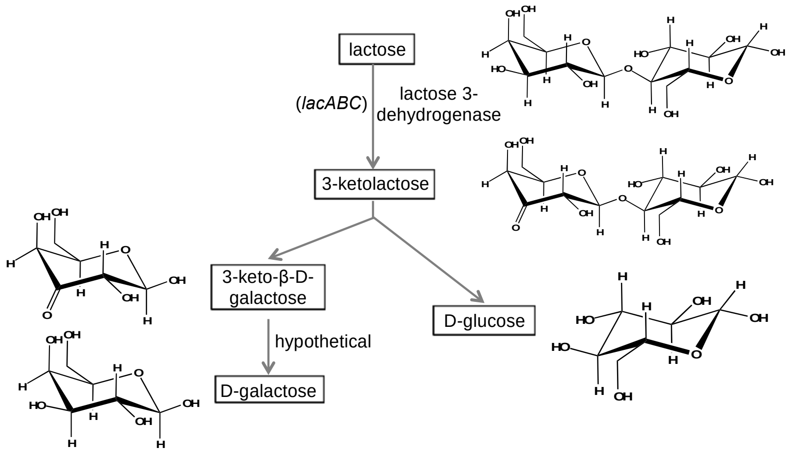


**Supplementary Figure 2. Putative 3-ketoglycoside pathways.** Proposed pathways (from MetaCyc) for (A) sucrose catabolism in *Agrobacterium tumefaciens* and (B) lactose catabolism in *Caulobacter crescentus*.

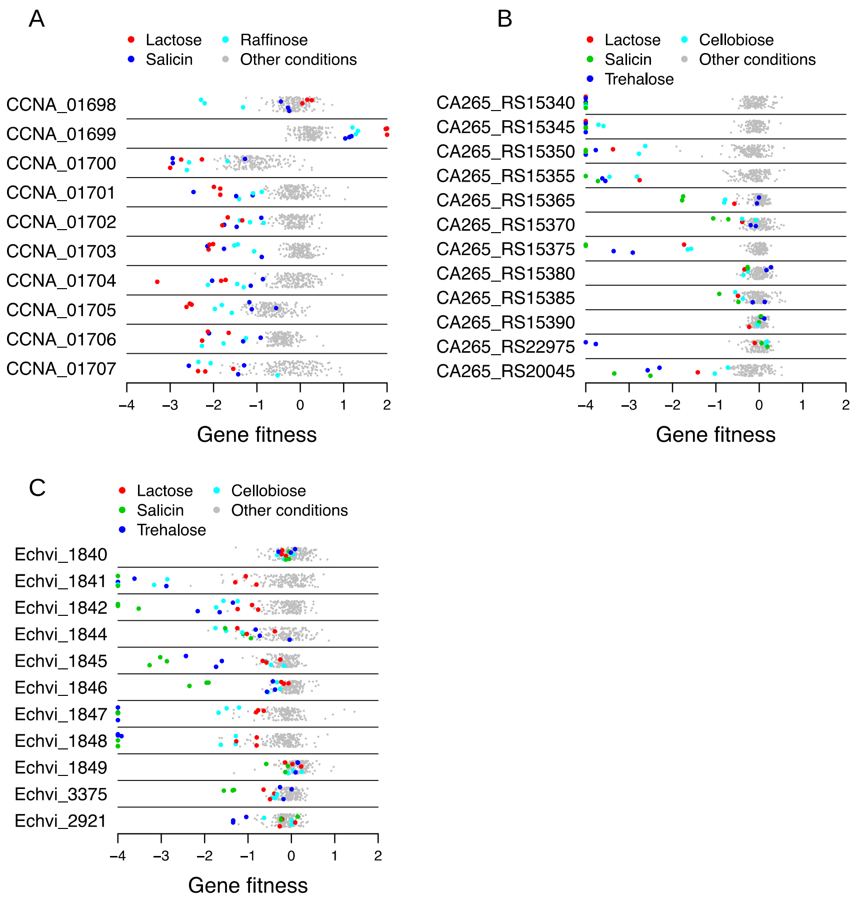


**Supplementary Figure 3. Fitness data for putative 3-ketoglycoside pathway genes in three bacteria.** Gene fitness data for select genes in (A) *Caulobacter crescentus*, (B) *Pedobacter* sp. GW460-11-11-14-LB5, and (C) *Echinicola vietnamensis* KMM6221. The *y*-axis is random and values < -4 are shown at -4.

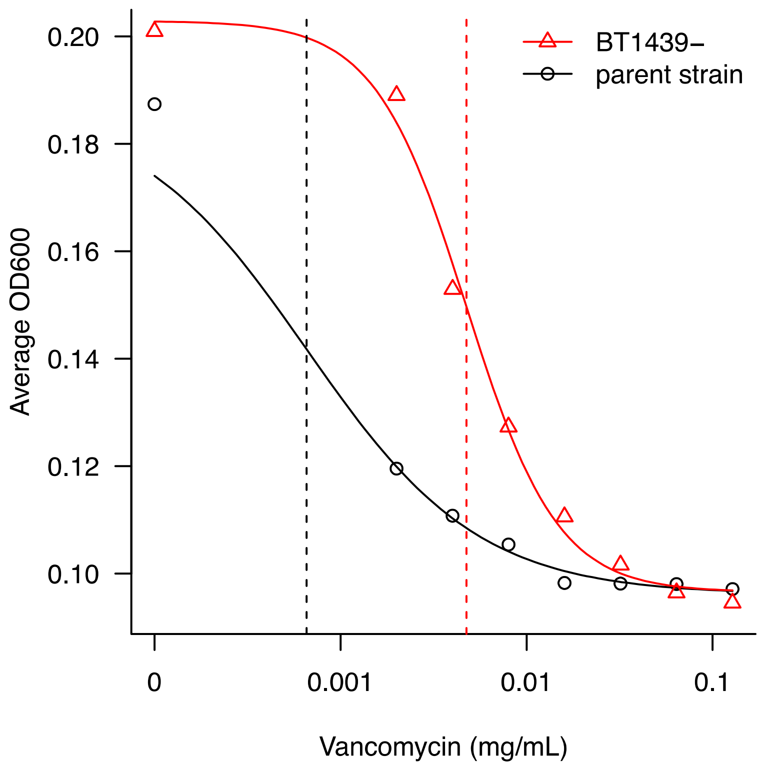


**Supplementary Figure 4. BT1439 is detrimental to fitness on vancomycin.** Dose-response curves for a BT1439 deletion mutant and the parental strain on vancomycin. Each point is the average of 3 replicate growth curves (for the mutant) or 4 replicate growth curves (for the parent strain). The vertical dotted lines indicate the $IC_{50}$ values. The curves are the fit as determined using the R package drc (Methods).
